# Flow-independent accumulation of motor-competent non-muscle myosin II in the contractile ring is essential for cytokinesis

**DOI:** 10.1101/333286

**Authors:** DS Osorio, FY Chan, J Saramago, J Leite, AM Silva, AF Sobral, R Gassmann, AX Carvalho

**Affiliations:** Instituto de Investigação e Inovação em Saúde – i3S, Universidade do Porto, Portugal.; Instituto de Biologia Molecular e Celular – IBMC, Porto, Portugal.

## Abstract

Cytokinesis in animal cells requires the assembly of a contractile actomyosin ring, whose subsequent constriction physically separates the two daughter cells. Non-muscle myosin II (myosin) is essential for cytokinesis, but the role of its motor activity remains poorly defined. Here, we examine cytokinesis in *C. elegans* one-cell embryos expressing myosin motor mutants generated by genome editing. Motor-dead myosin, which is capable of binding F-actin, does not support cytokinesis, and embryos co-expressing motor-dead and wild-type myosin are delayed in cytokinesis. Partially motor-impaired myosin also delays cytokinesis and renders contractile rings more sensitive to reduced myosin levels. Thus, myosin motor activity, rather than its ability to cross-link actin filaments, drives contractile ring assembly and constriction. We further demonstrate that myosin motor activity is required for long-range cortical actin flows, but that flows *per se* play a minor role in contractile ring assembly. Our results suggest that flow-independent recruitment of motor-competent myosin to the cell equator is both essential and rate-limiting for cytokinesis.

## Introduction

Cytokinesis is the final step of cell division that leads to the partitioning of the mother cell into two daughter cells, ensuring that each retains one copy of the replicated genome. Although cell-substrate adhesion may facilitate division in cultured cells (Neujahr et al., 1997; Kanada et al., 2005; Nagasaki et al., 2009; Dix et al., 2018), in fungi and animals cytokinesis primarily relies on acto-myosin dependent force generation and on the assembly and constriction of a distinct acto-myosin structure, the contractile ring, that forms at the cell equator. The contractile ring assembles beneath the plasma membrane after anaphase onset and subsequently constricts, folding the cell membrane inwards to achieve the physical separation between daughter cells (Green et al., 2012). The major components of the contractile ring are filaments of actin and non-muscle myosin II (hereafter myosin). Actin filaments polymerize and depolymerize during ring constriction (Carvalho et al., 2009; Stachowiak et al., 2014; Silva et al., 2016; Wollrab et al., 2016), and actin filament dynamics are controlled by a variety of actin-binding proteins (Blanchoin et al., 2014). Myosin is a hexameric protein complex composed of a dimer of heavy chains and two pairs of light chains. Each heavy chain has a N-terminal globular head that contains an ATP-binding pocket and an actin binding site, a lever arm where the light chains bind, and a C-terminal coiled-coil domain involved in interactions that promote heavy chain dimerization and formation of multi-headed bipolar filaments (Niederman and Pollard, 1975; Vicente-Manzanares et al., 2009). ATP hydrolysis induces coupled conformational changes that are transmitted through the head subdomains to the lever arm, generating a power stroke that causes myosin to move towards the actin filament barbed (+) end. In an interconnected actin-filament network with antiparallel filament arrangement, this movement causes actin filaments to slide past one another and the network to contract. In addition, the ability to bind actin allows myosin filaments to exert tension and maintain the network connected. Myosin is essential for cytokinesis in different systems (Mabuchi and Okuno, 1977; De Lozanne and Spudich, 1987; Straight et al., 2003), but the requirement for myosin motor activity has been a subject of debate. Budding yeast is able to perform cytokinesis in the presence of a motor-less myosin (Lord et al., 2005; Mendes Pinto et al., 2012). In fission yeast myosin motor activity appears to be required for cytokinesis, but additional myosins also contribute (Laplante et al., 2015; Laplante and Pollard, 2017; Palani et al., 2017). In *Dictyostelium* cells, specific mutations within the ATPase domain result in motor-dead myosins that when expressed in suspension cells cause growth phenotypes similar to or more severe than those of cells expressing no myosin, highlighting the importance of motor activity in this system (Shimada et al., 1997; Sasaki et al., 1998). In animals, whether myosin motor activity is absolutely required is less clear, since studies have relied on the use of the small molecule inhibitor blebbistatin, depletion/inactivation of myosin/myosin temperature-sensitive mutants, or non-phosphorylatable mutants of the regulatory light chain that is required for myosin complex activation (Straight et al., 2003; Davies et al., 2014; Reymann et al., 2016; Descovich et al., 2018). None of these approaches can provide a definitive answer regarding the requirement for motor activity, since blebbistatin locks myosin in a low actin-affinity state (Kovács et al., 2004), depletion or inactivation of myosin does not differentiate between motor and cross linking capacities of myosin, and interfering with the regulatory light chain may affect other myosins or influence myosin II localization and/or structure (Heissler and Sellers, 2015; Liu et al., 2016; Vasquez et al., 2014). The effect of specific motor-impairing mutations was reported for one of the mammalian myosin heavy chains in COS-7 cells and mouse cardiomyocytes (Ma et al., 2012). In this study, the requirement for myosin motor activity was contested, because apparently motor-dead mutants that still bind actin filaments were sufficient to replace wild-type myosin during cytokinesis.

Ultrastructural studies show that filamentous actin (F-actin) in the contractile ring consists primarily of unbranched filaments aligned parallel to the ring circumference and arranged in an antiparallel manner (Schroeder, 1973; Sanger and Sanger, 1980; Maupin and Pollard, 1986; Kamasaki et al., 2007; Henson et al., 2017). Additionally, myosin has been shown to form arrays of aligned filaments or stacks of filaments running parallel to actin filaments, an organization that is compatible with a purse-string mechanism where actin filament sliding by myosin would drive ring constriction (Fenix et al., 2016; Henson et al., 2017). Platinum replica transmission electron microscopy of isolated cortices of dividing sea urchin eggs revealed that the cortex outside of the contractile ring has a different network architecture, with actin filaments establishing a homogeneous isotropic meshwork with some dense foci of material (Schroeder, 1973; Sanger and Sanger, 1980; Maupin and Pollard, 1986; Kamasaki et al., 2007; Henson et al., 2017). How myosin distributes in the remaining cortex is less clear. In *C. elegans*, myosin appears in patches that flow toward the cell equator, and transient cortical myosin domains have been observed in NRK-52E epithelial cells (Werner et al., 2007; Zhou and Wang, 2008; Dickinson et al., 2013). Actin and myosin cortical flows towards the cell equator during early cytokinesis have been described in several experimental systems but are less evident in some mammalian cells and are absent in yeast, where myosin accumulates in nodes at the ring site (Cao and Wang, 1990; Yumura, 2001; Wu et al., 2006; Murthy and Wadsworth, 2005; Vavylonis et al., 2008; Laplante et al., 2015; Spira et al., 2017).

The *C. elegans* embryo is particularly suited for the quantitative *in vivo* analysis of cytokinesis, as embryos are large and its divisions stereotypical and temporally invariant. *C. elegans* possesses two non-muscle myosin II heavy chains, NMY-1 and NMY-2. NMY-2 has been shown to be essential for cytokinesis (Guo and Kemphues, 1996; Cuenca et al., 2003; Davies et al., 2014), whereas NMY-1 is required during late embryonic development (Piekny et al., 2003) and in the adult gonad and spermatheca (Kovacevic et al., 2013; Coffman et al., 2016; Wirshing and Cram, 2017). In this study we characterize NMY-2 mutants generated by genome editing to assess the role of motor activity during cytokinesis in the *C. elegans* early embryo. Our results suggest that it is myosin motor activity, and not myosin’s ability to cross-link actin filaments or modulate actin filament levels, that drives ring assembly and constriction. We show that while cortical actin flows depend on myosin motor activity, cytokinesis can complete in the absence of flows and that flow-independent equatorial accumulation of motor-competent myosin determines the timing of contractile ring assembly and constriction initiation.

## Results

### Generation of *C. elegans* motor-dead non-muscle myosin II mutants that bind F-actin

A previous alanine mutagenesis screen in the highly conserved switch I region of the ATPase domain of *D. discoideum* non-muscle myosin II yielded a series of mutants with compromised motor activity (Shimada et al., 1997). We sought to take advantage of the high sequence conservation among myosins to generate equivalent mutants in *C. elegans*. We chose two point mutations, corresponding to S251A and R252A in NMY-2, that were shown to yield motor-dead myosin in *D. discoideum* (Fig. 1A, B, S1A). To assess the potential of these mutations to affect myosin motor activity in *C. elegans,* we first examined the consequences of introducing the mutations into muscle myosin. Muscle fibers are composed of sarcomeres, which require myosin motor activity to contract. Although muscle and non-muscle myosin II present differences in ATP hydrolysis kinetics and motility rates, the principle underlying the change in molecular conformation that allows for the power stroke necessary to translocate actin filaments is identical in both motors and relies on extremely well conserved regions, including the Switch I loop of the ATP binding site (Heissler and Sellers, 2016, Fig. 1C, S1A). Previous studies have established that the mechanistic effects of mutations in conserved myosin head residues are transposable between class II myosins and even between different myosin classes (Li et al., 1998; Onishi et al., 1998a; Forgacs et al., 2009; Trivedi et al., 2012). Using CRISPR-Cas9-based genome editing (Arribere et al., 2014), we introduced the two point mutations into UNC-54, the main skeletal muscle myosin that is essential for motility and egg laying (Epstein and Thomson, 1974; Fire et al., 1991). We were able to generate homozygous animals expressing UNC-54(S240A), corresponding to NMY-2(S251A), and UNC-54(R241A), corresponding to NMY-2(R252A). To evaluate muscle function, we monitored worm locomotion and egg laying rates (Fig. 1D, E). *Unc-54(S240A)* and *unc-54(R241A)* animals displayed a drastic reduction in liquid locomotion (0.2±0.1 Hz vs 1.6±0.1 Hz in controls) and were unable to lay eggs, as expected for strongly motor-impaired myosins. Residual movement observed in *unc-54(S240A)* and *unc-54(R241A)* animals was due to the secondary body wall muscle myosin MYO-3, since depletion of MYO-3 in either *unc-54* mutant led to paralysis on food plates and complete loss of motility in liquid (Fig. 1D). Complete loss of motility was also observed when UNC-54 and MYO-3 were co-depleted by RNAi in worms expressing wild-type UNC-54 (Fig. S1B). Interestingly, phalloidin staining of muscles in *unc-54(S240A)* and *unc-54(R241A)* animals revealed that actin organization in sarcomeres was preserved (Fig. S1C). This suggests that myosin motor activity is not required for actin organization in adult muscles, which agrees with recent findings in *D. melanogaster* (Loison et al., 2018). We conclude that mutating the highly conserved residues S240 and R241 results in non-functional UNC-54 *in vivo*, in agreement with the effects of the mutations on myosin motor activity reported *in vitro* for *D. discoideum* non-muscle myosin II. Of note, we also tested the corresponding mutation of R709C in mammalian myosin IIB (Fig. S2A), a disease-related mutation on the SH1 helix of the myosin molecule (Hu et al., 2002; Shibata et al., 2017), which was previously reported to be unable to translocate actin filaments (Ma et al., 2012). We found that UNC-54(R710C) only partially impaired muscle function. UNC-54(R710C) is therefore unlikely to be motor-dead (Fig. S2B,C).

**Figure 1.**
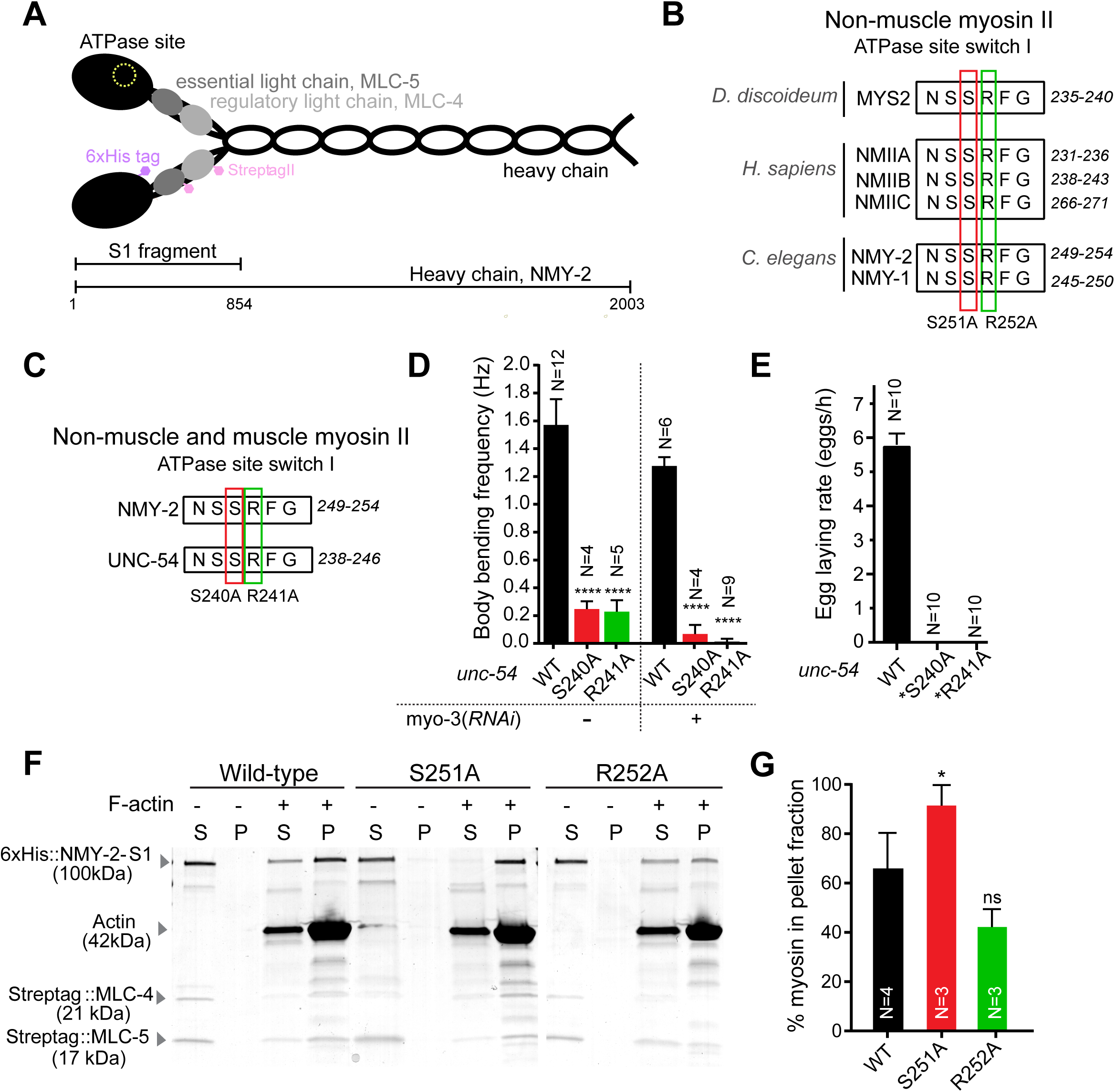
Establishing motor dead non-muscle myosin II mutants in *C. elegans*. **(A)** Schematic of the non-muscle myosin II hexamer indicating the S1 fragment used for actin interaction assays. **(B)** Alignment of *D. discoideum*, *H. sapiens* and *C. elegans* non-muscle myosin IIs showing the conservation of the ATPase switch I domain. Two mutated residues tested in this study are in red and green boxes (S251A, R252A, respectively, numbered as in *C. elegans* NMY-2). **(C)** Switch I residues are also conserved in UNC-54, the major body-wall muscle myosin heavy chain in *C. elegans.* **(D)** Body bend frequency in liquid ± 95% CI in wild-type and *unc-54* motor mutant animals depleted or not of MYO-3, the secondary body wall muscle myosin. **(E)** Egg laying rate ± 95% CI in wild-type and *unc-54* mutant animals. *unc-54(S240A)* and *unc-54(R241A)* animals do not lay eggs but embryos are viable and develop normally inside the mother (*). **(F)** Coomassie stained SDS-PAGE gel of high-speed F-actin cosedimentation assays where NMY-2 S1 fragments carrying the indicated mutations were incubated with or without 14.7 μM of F-Actin before being submitted to ultracentrifugation and separated in supernatant (S) and pellet (P) fractions. **(G)** Percentage ± 95% CI of NMY-2 S1 fragment present in the pellet. N is the number of worms analyzed in (D) and (E) and the number of independent experiments in (G). Statistical significance was determined using one-way ANOVA followed by Bonferroni’s multiple comparison test; * P<0.01, **** P<0.00001, ns = not significant (P>0.05).

Next, we assessed the ability of NMY-2(S251A) and NMY-2(R252A) to bind F-actin in high speed co-sedimentation assays. His-tagged NMY-2 S1-like fragments (residues 1-854; routinely used for actin binding and kinetic assays; Manstein et al., 1989) carrying either mutation were expressed in Sf21 insect cells along with the myosin regulatory (MLC-4) and essential (MLC-5) light chains that were N-terminally tagged with Strep-tag II. NMY-2(S1)-MLC4-MLC5 complexes were purified and incubated with F-actin before high-speed centrifugation. In the control experiments, where no F-actin was added, all myosin was present in the supernatant, indicating that wild-type and mutant S1 versions were equally soluble. Conversely, all S1 versions were found in the pellet along with F-actin, showing that the mutants are capable of binding actin filaments (Fig. 1F). Quantification of the protein fraction present in the supernatant and pellet suggested that the S251A mutant had a higher affinity for F-actin (91±11 vs 66±23% for wild-type S1), whereas the distribution of the R252A mutant was similar to wild-type S1 (42±18 vs 66±23% for wild-type S1, p=0.0641; Fig. 1G). We conclude that NMY-2(S251A) and NMY-2(R252A) represent two motor-dead myosin mutants that are able to bind F-actin.

### Myosin motor activity is essential for embryo production and development

Next, we generated animals expressing NMY-2(S251A) or NMY-2(R252A) using genome editing (Arribere et al., 2014). Animals homozygous for either mutation exhibited severe gonad malformation and were consequently sterile (Fig. 2A). To examine the impact of these mutants on embryogenesis, we introduced transgene-encoded wild-type NMY-2::mCherry into *nmy-2(S251A)* and *nmy-2(R252A)* animals. NMY-2::mCherry was expressed from the *nmy-2* promoter and 3’UTR and the transgene was partially re-encoded so it could be specifically depleted by RNAi (NMY-2::mCherry^sen^, sen indicating RNAi-sensitive; Fig. 2B). The resulting strains were homozygous for both versions of NMY-2, which were expressed at identical levels (Fig. 2C). The presence of NMY-2::mCherry^sen^ allowed homozygous *nmy-2(S251A)* and *nmy-2(R252A)* mutants to develop normally and lay eggs. However, when NMY-2::mCherry^sen^ was depleted, *nmy-2(S251A)* and *nmy-2(R252A)* embryos were not viable, demonstrating that motor-dead NMY-2 does not support embryonic development (Fig. 2D).

**Figure 2.**
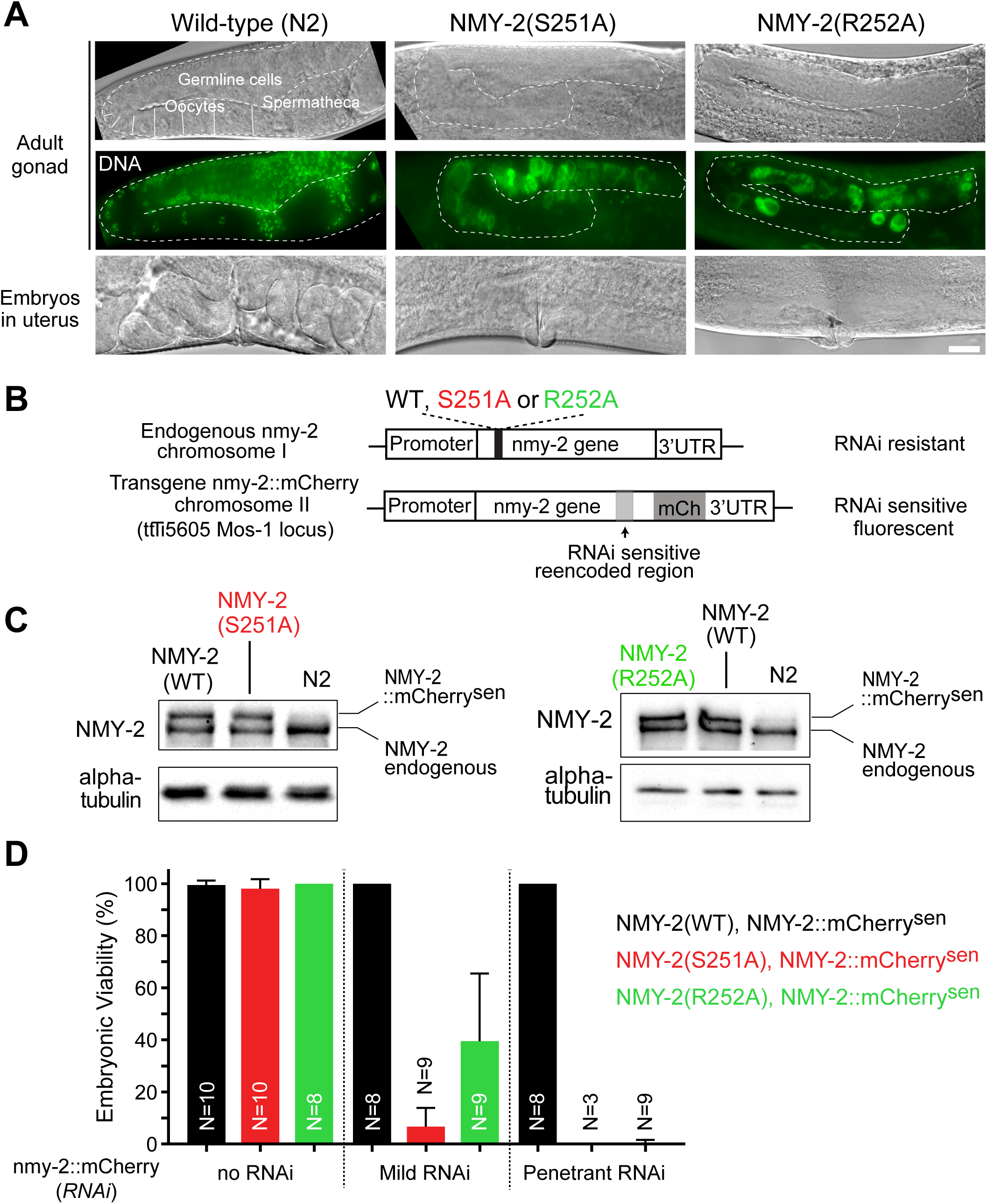
Motor-dead myosins do not support embryonic development. **(A)** Differential interference contrast images of the gonad (top) or uterus (bottom) in wild type, *nmy-2(S251A)/nmy-2(S251A)* or *nmy-2(R252A)/nmy-2(R252A)* mutant adult animals. Fluorescence images of DAPI labelled DNA in the gonad region is shown in the central row. **(B)** Schematic of the endogenous and transgenic *nmy-2* loci. Mutations were introduced in the endogenous nmy-2 gene in chromosome I by CRISPR/Cas9. A wild-type transgenic version of nmy-2 carrying a re-encoded portion for RNAi sensitivity and fused to mCherry (NMY-2::mCherry^sen^) was introduced in single copy in a defined position of chromosome II using MosSCI. **(C)** Immunoblot showing protein levels of endogenous and transgene-encoded NMY-2::mCherry^sen^ in wild-type and mutant animals. **(D)** Embryonic viability ± 95% CI of the progeny of (N) animals of the strains shown in (B) when NMY-2::mCherry^sen^ is depleted or not by RNAi. Short-term and long-term RNAi conditions were used to achieve mild or penetrant protein depletion. Scale bars, 10 μm.

### Motor-dead myosins do not support cytokinesis

Next we asked whether NMY-2(S251A) and NMY-2(R252A) could support cytokinesis in dividing one-cell embryos. Expression of the motor-dead myosins did not prevent wild-type NMY-2::mCherry^sen^ from localizing in cortical patches. However, myosin patches outside the cell equator were less abundant than in controls (Fig. 3A, Supplemental Video S1). Interestingly, cytokinesis was slightly delayed in *nmy-2(S251A)* and *nmy-2(R252A)* mutants, suggesting that the presence of motor-dead myosin interfered with the function of NMY-2::mCherry^sen^ (Fig. 3C). Specific depletion of NMY-2::mCherry^sen^ by RNAi did not alter cytokinesis kinetics in embryos expressing wild-type endogenous NMY-2. By contrast, penetrant depletion of NMY-2::mCherry^sen^ in *nmy-2(S251A)* or *nmy-2(R252A)* embryos always caused cytokinesis failure. When NMY-2::mCherry^sen^ was mildly depleted using shorter RNAi treatment, some embryos still failed cytokinesis, while the majority of embryos completed cytokinesis but did so with slower kinetics than non-depleted embryos (Fig. 3B). To assess cytokinesis kinetics in more detail, we measured 1) the time interval between anaphase onset and the formation of a shallow equatorial deformation, when contractile ring components are being recruited to the cell equator (“Ring Assembly”), 2) the time interval between shallow deformation and the folding of the plasma membrane into a back-to-back configuration (“Furrow Initiation”), and 3) the rate of ring constriction (Fig. 3B’). Abscission in *C. elegans* embryos only completes in the following round of cell divisions and was not analyzed (Green et al., 2013). Mild depletion of NMY-2::mCherry^sen^ in embryos expressing motor-dead myosins delayed all stages of cytokinesis (Fig. 3C): time for ring assembly and furrow initiation was increased (ring assembly: 62±6 s for controls, 219±26 s for S251A, 201±30 s for R252A; furrow initiation: 59±8 s for controls, 136±19 s for S251A and 128±17 s for R252A), and ring constriction rate was reduced (0.18±0.01 μm/s for controls, 0.11±0.01 μm/s for S251A and 0.12±0.01 μm/s for R252A). Moreover, the remaining NMY-2::mCherry^sen^ accumulated slowly and solely at the equatorial region until furrowing initiated (Supplemental Video S4). Together, these data show that expression of motor-dead myosin disrupts the localization of wild-type myosin outside the equator and that cytokinesis is progressively affected as the ratio of motor-dead myosin to wild-type myosin increases. We conclude that myosin motor activity is required for ring assembly, furrow initiation and ring constriction and that its ability to crosslink F-actin is not sufficient for cytokinesis.

**Figure 3.**
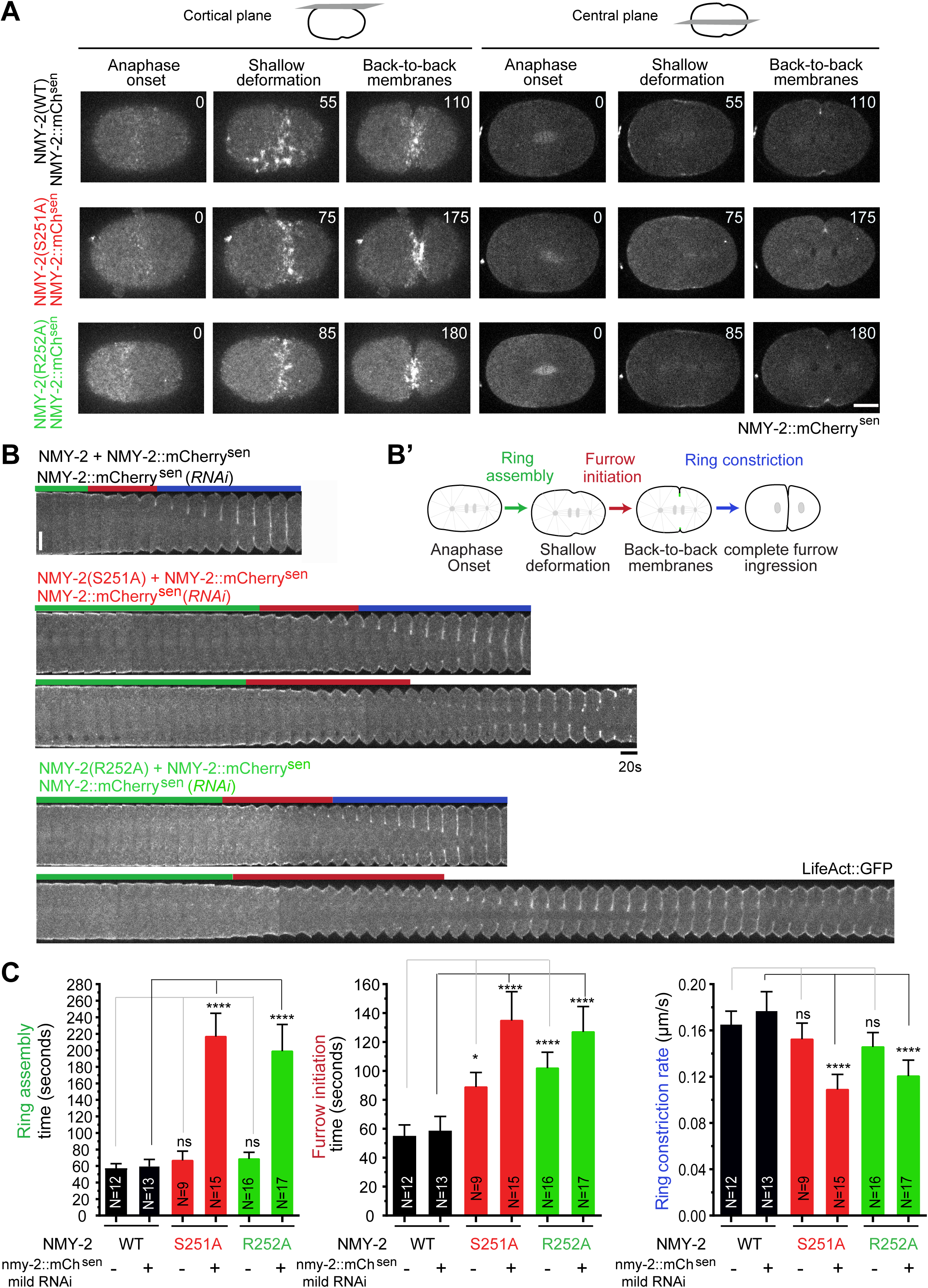
Myosin motor activity is essential for cytokinesis. **(A)** Stills from time-lapse imaging series of the cortical or central planes in embryos expressing NMY-2(WT), NMY-2(S251A) or NMY-2(R252A), along with NMY-2::mCherry^sen^. **(B, B’)** Kymographs of the equatorial region of embryos co-expressing LifeAct::GFP and wild-type or mutant NMY-2 along with NMY-2::mCherry^sen^ after mild depletion of NMY-2::mCherry^sen^ (B). Examples of mutant embryos that failed cytokinesis using the same RNAi conditions are shown on the corresponding bottom rows. Eight out of 23 *nmy-2(S251A)* and 3 out of 19 *nmy-2(R252A)* embryos failed cytokinesis. First frame corresponds to anaphase onset. Green, red and blue bars indicate the intervals of equatorial band formation, furrow initiation and ring constriction as depicted in (B’). **(C)** Mean duration ± 95% CI of ring assembly and furrow initiation, as indicated in (B’), and ring constriction rate in wild-type or mutant one-cell embryos subjected or not to mild depletion of NMY-2::mCherry^sen^. N is the number of embryos analyzed. Statistical significance was determined using one-way ANOVA followed by Bonferroni’s multiple comparison test; * P<0.01, **** P<0.00001, ns = not significant (P>0.05). Scale bars, 10 μm.

### Partial impairment of myosin motor activity slows ring constriction and reduces the robustness of cytokinesis

Next, we generated animals expressing NMY-2(S250A), which is predicted to result in partially motor-compromised myosin, as the corresponding mutant in *D. discoideum* exhibited reduced motor activity *in vitro* (Shimada et al., 1997). Like S251 and R252, the S250 residue is located in the switch I region of the ATPase domain of myosin (Fig. 1B, S1A). We were able to generate homozygous animals expressing the equivalent mutation in muscle myosin, *unc-54(S239A)* (Fig. 1C). These animals displayed reduced locomotion in liquid (body bends swimming frequency of 0.6±0.1 Hz vs 1.6±0.1 Hz in controls), and reduced egg laying rate (3.7±0.5 vs 5.8±0.3 eggs per hour in controls), as expected for a myosin with partially-impaired motor activity (Fig. 4A, B). The S1 fragment of NMY-2(S250A) co-sedimented with F-actin similarly to the wild-type counterpart (Fig. 4C, D). We were also able to generate homozygous animals expressing NMY-2(S250A) at levels comparable to wild-type controls (Fig. 4E, F). These animals were fully viable and propagated normally (Fig. 4G). To characterize cytokinesis in early embryos, we introduced LifeAct::GFP into the *nmy-2(S250A)* mutants. Embryos expressing NMY-2(S250A) had normal ring assembly times (51±5 s for S250A, 56±4 s for controls) but displayed increased furrow initiation times (79±8 s for S250A, 59±8 s for controls) and reduced ring constriction rates (0.12±0.004 μm/s for S250A, 0.17±0.01 μm/s for controls; Fig. 4H, I). Thus, exclusive expression of partially compromised motor myosin has the same effect on the kinetics of cytokinesis as co-expressing similar amounts of motor-dead and wild-type myosin (compare Fig. 3C, no RNAi, with Fig. 4I). We also generated homozygous worms expressing NMY-2(R718C), which corresponds to R709C in mammalian myosin IIB (Fig. S2A, D-G). Characterization of cytokinesis revealed delays that were similar to those we observed in embryos expressing NMY-2(S250A) (Fig. S2G). Together with the partial effect of the equivalent mutation in UNC-54 on muscle function (Fig. S2B, C) these results are consistent with the idea that this mutation compromises, but does not abolish, myosin motor activity.

**Figure 4.**
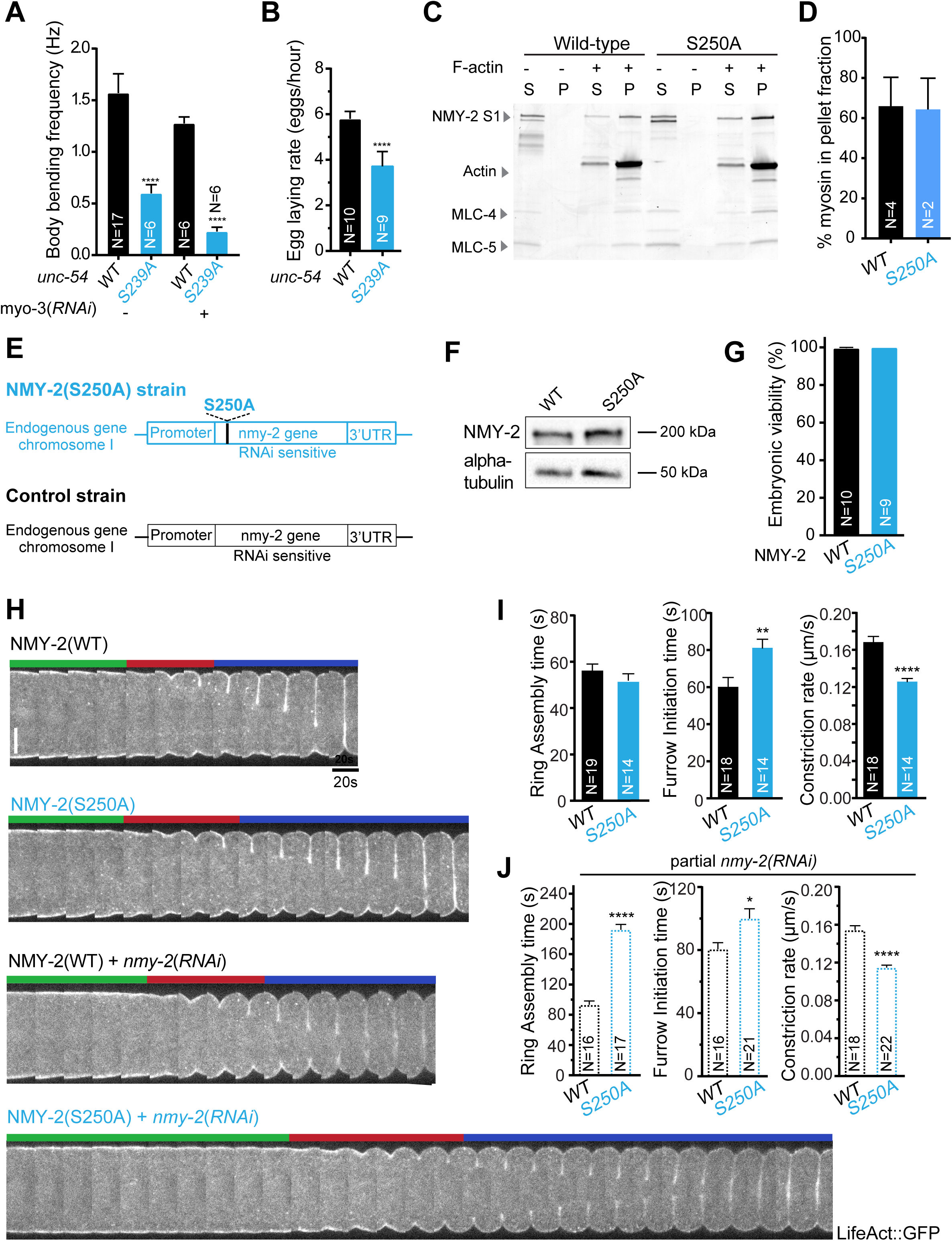
Partial impairment of myosin motor activity slows ring constriction and reduces cytokinesis robustness. **(A)** Body bend frequency in liquid ± 95% CI in wild-type and *unc-54(S239A)* mutant animals depleted or not of MYO-3, the secondary body wall muscle myosin. **(B)** Egg laying rate ± 95% CI in wild-type and *unc-54(S239A)* animals. **(C)** Coomassie stained SDS-PAGE gel of high-speed F-actin co-sedimentation assays where wild-type or S250A NMY-2 S1 fragments were incubated with or without 14.7 μM of F-Actin before being submitted to ultracentrifugation. (S) indicates the supernatant and (P) the pellet fractions. **(D)** Percentage ± 95% CI of NMY-2 S1 fragment present in the pellet. **(E)** Schematic of the *nmy-2* locus after introduction of the S250A mutation by CRISPR/Cas9. **(F)** Immunoblot showing protein levels of wild-type or NMY-2(S250A) endogenous protein in wild-type or mutant animals, respectively. Alpha-tubulin is used as loading control. **(G)** Embryonic viability ± 95% CI of progeny of (N) wild-type or *nmy-2(S250A)* mutant animals. **(H)** Kymographs of the equatorial region of embryos co-expressing LifeAct::GFP and wild-type or mutant NMY-2. Last 2 rows correspond to wild-type or mutant embryo after mild depletion of NMY-2. First frame corresponds to anaphase onset. Green, red and blue bars indicate the intervals of equatorial band formation, furrow initiation and ring constriction as depicted in figure 3B’. **(I, J)** Mean duration ± 95% CI of ring assembly and furrow initiation, and ring constriction rate in wild-type or mutant one-cell embryos subjected or not to mild depletion of NMY-2. N is the number of independent experiments in (D), the number of worms whose progeny was analyzed in (G) and the number of analyzed embryos in (I, J). Statistical significance was determined using one-way ANOVA followed by Bonferroni’s multiple comparison test; * P<0.01, ** P<0.001, **** P<0.00001. Scale bar, 10 μm.

Next, we asked whether embryos expressing wild-type or NMY-2(S250A) were equally sensitive to a decrease in overall myosin levels. In mild RNAi conditions (short course of RNAi treatment), embryos expressing wild-type myosin always completed cytokinesis and exhibited mild delays in ring assembly (93±11 s), furrow initiation (81±9 s) and constriction rate (0.15±0.01 μm/s; Fig. 4J). In contrast, mild depletion of NMY-2 in embryos expressing NMY-2(S250A) led to a substantial increase in ring assembly and furrow initiation times (192±15 s and 100±13 s, respectively) and had a stronger effect on ring constriction rate (0.11±0.01 μm/s, Fig. 4J). Additionally, 4 out of 22 NMY-2(S250A)-expressing embryos depleted of NMY-2 failed cytokinesis. We conclude that contractile rings assembled with motor-compromised myosin are less resilient to a decrease in myosin levels than rings assembled with wild-type NMY-2.

### Perturbation of myosin levels or motor activity does not change actin levels during ring constriction

Next, we examined whether myosin modulated actin levels in the contractile ring. Interestingly, we found that the amount of actin in the constricting ring at 50% ingression did not vary in one-cell embryos expressing NMY-2(S250A), or in embryos expressing NMY-2(S251A) or NMY-2(R252A) in the presence of normal or decreased levels of NMY-2::mCherry^sen^ (Fig. 5A). The same was observed in embryos partially depleted of NMY-2 (Fig. 5A). In addition, just like in control embryos, the concentration of LifeAct::GFP in the contractile ring did not change throughout constriction when myosin was partially depleted (Fig. 5B), indicating normal net depolymerization of actin filaments. We conclude that neither lowering myosin levels nor inhibiting myosin motor activity affects actin levels in the constricting ring.

**Figure 5.**
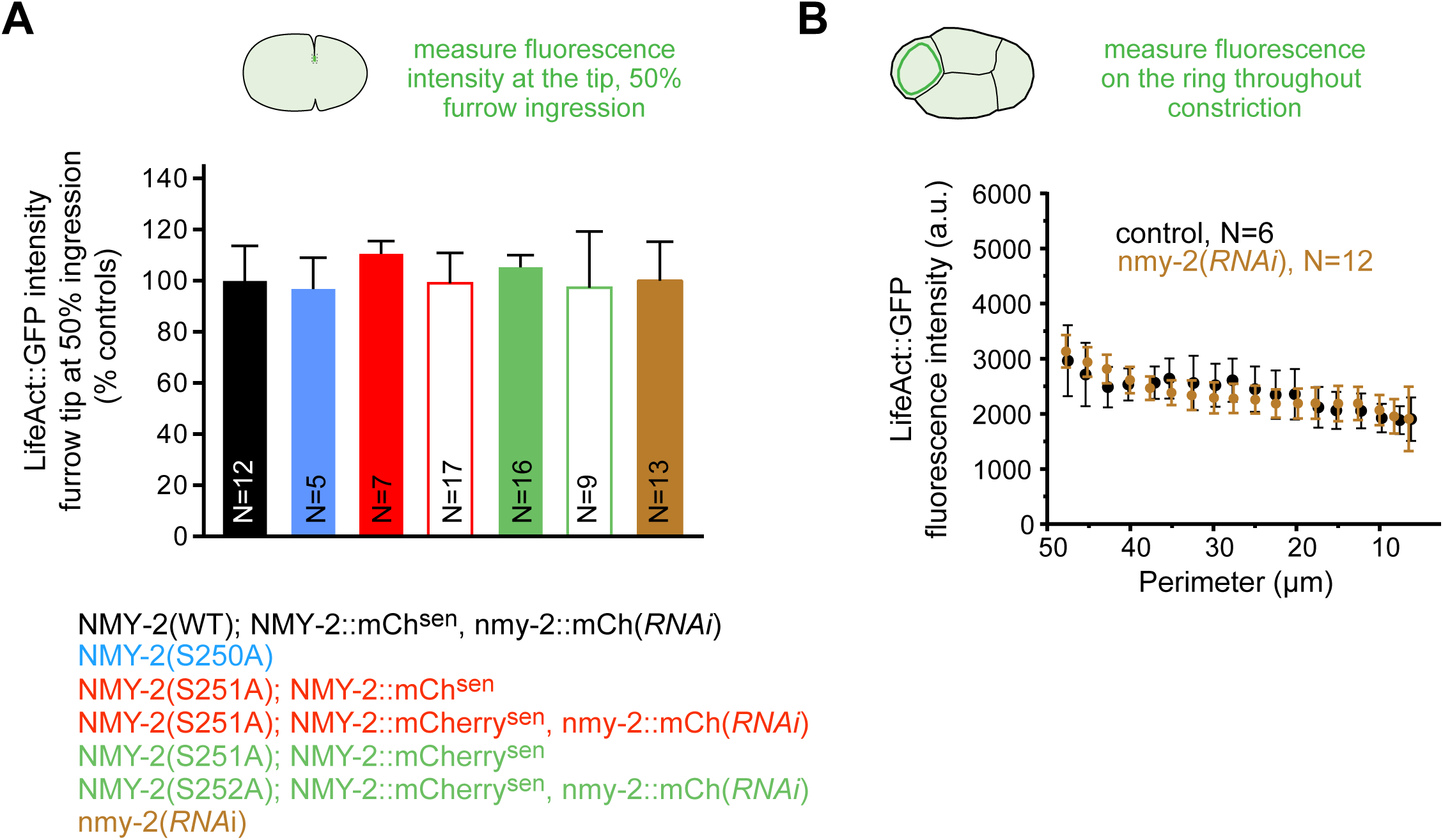
Myosin motor activity does not affect actin levels in the contractile ring during constriction. **(A)** LifeAct::GFP levels ± 95% CI in the contractile ring at 50% furrow ingression (schematic on top) in control embryos, embryos expressing NMY-2(S250A), and embryos co-expressing NMY-2(S251A) or NMY-2(R252A) and NMY-2::mCherry^sen^ with or without mild depletion of NMY-2::mCherry^sen^. Values were normalized to corresponding controls. **(B)** Quantification of LifeAct::GFP levels ± 95% CI during ring constriction in ABa cells of 4-cell embryos, where the ring can be observed end-on (schematic on top).

### Long-range cortical actin flows are dispensable for contractile ring formation and furrow ingression

A recent analysis in one-cell embryos expressing LifeAct::mKate2 suggested that alignment and compaction of actin filaments at the cell equator is driven by myosin-dependent unidirectional persistent cortical flows, and that flows are consequently important for cytokinetic furrow formation (Reymann et al., 2016). We therefore sought to evaluate the importance of myosin motor activity for cortical actin flows. As it has been reported that LifeAct expression has dose-dependent effects on actin dynamics and at high concentrations affects cytokinesis in yeast cells (Courtemanche et al., 2016), we first compared cytokinesis kinetics and cortical distribution of F-actin in embryos expressing LifeAct::mKate2, which was used in a previous study (Reymann et al., 2016), and in embryos expressing LifeAct::GFP (Silva et al., 2016). We found that in one-cell embryos expressing LifeAct::mKate2 furrow initiation was delayed and ring constriction rate was significantly reduced compared to controls (Fig. S4A). By contrast, the kinetics of cytokinesis in early embryos was unaffected in animals expressing LifeAct::GFP. Moreover, the cortical distribution of LifeAct::GFP, but not that of LifeAct::mKate2, was similar to that observed in embryos expressing endogenous fluorescently tagged Plastin^PLST-1^, a crosslinker that binds all F-actin pools during cytokinesis (Ding et al., 2017; Fig. S4B). Based on these results, we chose LifeAct::GFP for the analysis of cortical actin flows. We were able to monitor cortical actin flows in all myosin motor mutants, as well as in NMY-2 depleted embryos, because levels of cortical actin were unaffected (Fig. 3B, 4H, 5, 6A’, 7B, Supplemental Videos 2 and 4). Kymographs of the cortical region from anaphase onset to back-to-back membrane configuration revealed that actin flows directed from the posterior to the anterior were obvious in control embryos, were much less pronounced in embryos expressing NMY-2(S250A), and were absent in embryos expressing NMY-2(S251A) or NMY-2(R252A) with or without RNAi of NMY-2::mCherry^sen^ (Fig. 6A, 6A’, S5A). Flows were also not detected in embryos partially depleted of NMY-2 using the RNAi conditions shown in figure S3. Particle Image Velocimetry (PIV) analysis was performed to analyze cortical movement in more detail. The magnitude of velocity fields was calculated and color-encoded using a thermal scale in an arrow diagram where vector directionality is depicted (Fig. 6B, B’, Supplemental Video 2). Flow velocity was calculated by averaging the X-and Y-components (anterior-posterior and dorsal-ventral, respectively) of the velocity vectors in two regions positioned over the anterior or the posterior halves of the embryo (Fig. 6B, C). Average flow velocity was calculated for each time point between anaphase onset and back-to-back membrane configuration (Fig. 6C). In control embryos, anterior-directed cortical flows with a strong Y-component started just prior to equatorial shallow deformation, peaked shortly thereafter, and remained high until ∼50 seconds later (Supplemental Video 2). The direction of the flows was the same in the anterior and posterior sides of the equator, indicating that unlike during pseudocleavage (a transient furrow that forms in the early zygote during polarity establishment), counter-rotating flows do not occur to facilitate the initiation of cytokinesis (Fig. S5B; Naganathan et al., 2014). Interestingly, we found that the strong Y-component of the vectors was due to embryo compression, as the Y-component was drastically reduced in embryos imaged in open chambers, which completed cytokinesis with similar kinetics (Fig. S5C, D, Supplemental Video 3). In embryos solely expressing NMY-2(S250A), flows were observed for a shorter period of time and were of lower magnitude than in controls. Also distinct from controls, flows in NMY-2(S250A) diminished in intensity and became more erratic shortly after shallow deformation (Fig. 6B’, C, Supplemental Video 2). In striking contrast to controls, most vectors detected in embryos co-expressing NMY-2(S251A) or NMY-2(R252A) and NMY-2::mCherry^sen^ were extremely weak, had no coherent directionality, and did not increase in magnitude during the period when the ring formed and the furrow initiated ingression (Fig. 6B’, C, Supplemental Video 2). This profile was observed whether or not NMY-2::mCherry^sen^ was depleted by RNAi (Fig. S5E). We conclude that persistent and compressive actin cortical flows depend on motor-competent myosin and that in the absence of cortical flows contractile rings can form, furrows can continuously ingress and cytokinesis can complete.

**Figure 6.**
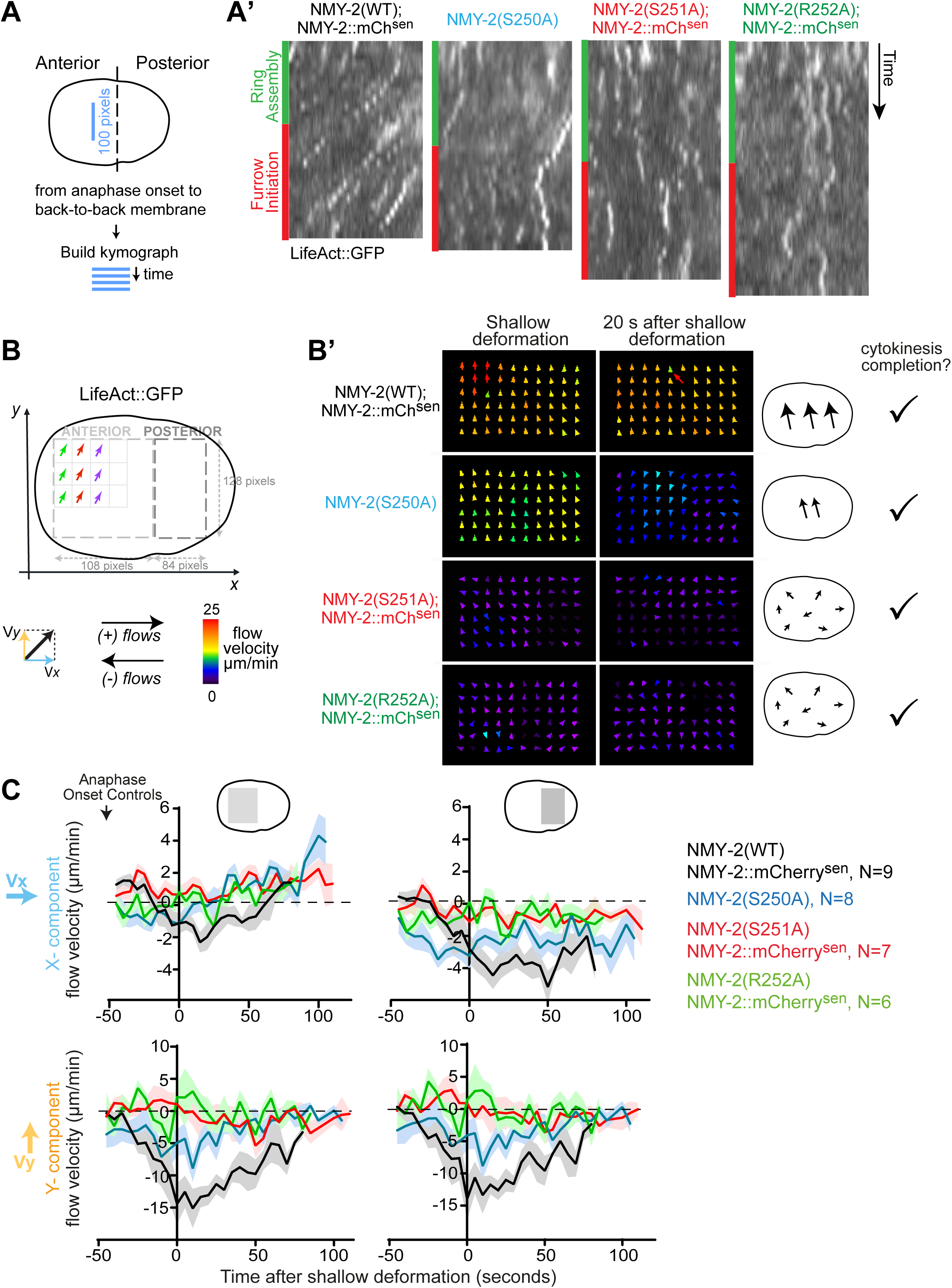
Long-range cortical actin flows are largely dispensable for contractile ring formation and furrow ingression. **(A)** Schematic illustrating region of the embryo with which kymographs in (A’) were built. **(A’)** Kymographs of the region shown in (A) in embryos co-expressing LifeAct::GFP and NMY-2::mCherry^sen^ as indicated. Green and red bars correspond to ring assembly and furrow initiation time intervals, as in figure 3B’. **(B)** Schematic illustrating how cortical LifeAct::GFP flows were analyzed by particle image velocimetry (PIV). **(B’)** Arrow diagrams depicting LifeAct::GFP flows in embryos of the indicated genotypes at shallow deformation and 20 seconds later (also see Supplemental Movie S2). Arrows represent average directionality and magnitude of vectors in each interrogation window. Vector magnitude is color-coded using a thermal scale as shown in B. The schematic on the right depicts overall flow intensity and directionality in each condition. **(C)** Average flow velocity LifeAct::GFP levels ± 95% CI in the anterior or posterior sides of embryos with indicated genotypes. All curves were aligned to the time of shallow deformation. X-component of the velocity vectors is shown on the top graphs and Y-component on the bottom graphs. N is the number of embryos analyzed.

### Equatorial accumulation of motor-competent myosin is rate limiting for contractile ring assembly and constriction initiation

The fact that equatorial actin bundles look relatively well aligned and organized in embryos expressing NMY-2(S250A) and in embryos co-expressing NMY-2(S251A) or NMY-2(R252A) and NMY-2::mCherry^sen^ (Supplemental Video 2) indicates that a flow-independent equatorial pool of myosin is sufficient for proper bundle alignment. We therefore more carefully examined actin bundle behavior and myosin recruitment during contractile ring assembly in embryos expressing myosin motor-dead mutants and in embryos depleted of NMY-2 (Supplemental Video 4; Fig. 7A). In control embryos at anaphase onset, cortical actin is enriched in puncta on an anterior cap and in an underlying actin meshwork that extends over the entire embryo. The first appearance of actin in the equatorial region (Actin Equatorial Appearance, Actin EA) is observed 30±7 s after anaphase onset (AO). A well-defined equatorial band (Actin EB) is visible shortly after (Actin EA-EB 20±4 s) and shallow equatorial deformation (SD) immediately follows (Actin EB-SD 12±3 s), coincident with the appearance of circumferential actin bundles running parallel to the division plane. Embryos expressing NMY-2(S251A) or NMY-2(R252A) after mild depletion of NMY-2::mCherry^sen^ displayed actin puncta distributed over the entire embryo at anaphase onset. This possibly reflects loss of polarity as a similar pattern of actin distribution is observed when PAR-2 is depleted (Fig. S4C). Actin equatorial enrichment is delayed relative to controls (Actin AO-EA 81±18 s and 98±28 s, respectively) and formation of an actin equatorial band is detected soon after (Actin EA-EB 26±4 s and 25±5 s, respectively). Equatorial actin bundles are slanted relative to the division plane and persist until shallow deformation eventually occurs (Actin EB-SD 136±48 s and 154±35 s, respectively; Fig. 7B). Polarity loss could explain the slight delay in actin equatorial appearance but not the extended interval between band formation and shallow deformation (Fig. S4D). Embryos partially depleted of NMY-2, displayed similar delays in actin equatorial appearance (Actin AO-EA 64±24 s), formation of the actin equatorial band (Actin EA-EB 25±5 s) and shallow deformation (Actin EB-SD 188±36 s). Equatorial actin bundles also looked slanted and also persisted until shallow deformation. We conclude that a minimal amount of motor-competent myosin needs to be present at the cell equator for proper actin bundle alignment and timely actin band formation and compaction.

**Figure 7.**
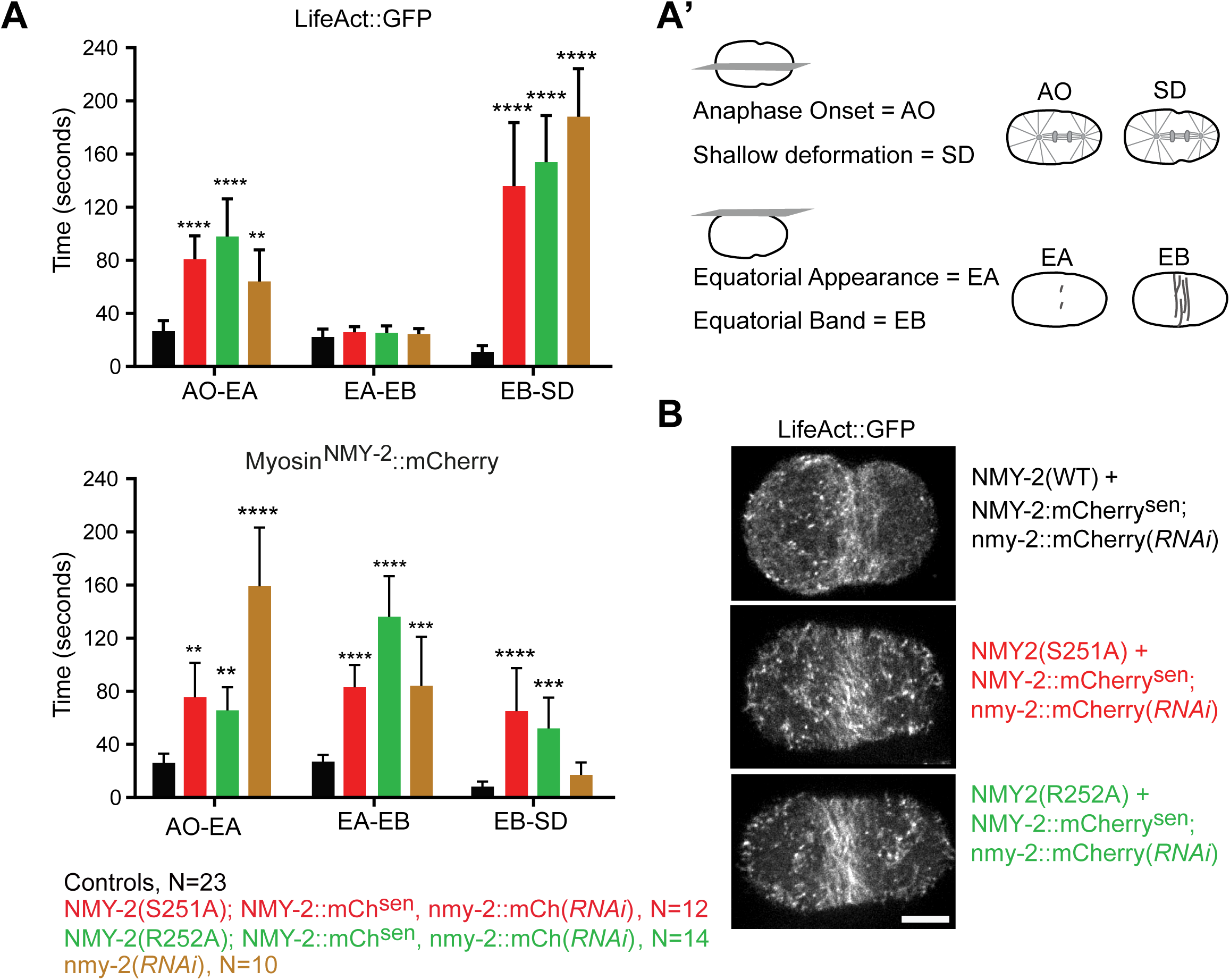
Flow-independent equatorial accumulation of enough motor-competent myosin is essential for cytokinesis. **(A, A’)** The interval of ring assembly was divided in 3 sub-intervals: Anaphase onset (AO) to first equatorial appearance of LifeAct::GFP or NMY-2::mCherry^sen^ (EA), EA to equatorial band formation (EB) of LifeAct::GFP or NMY-2::mCherry^sen^, and EB to shallow deformation (SD). AO and SD were determined by filming the embryo central plane and EA and EB by filming the embryo cortex (A’). The mean duration ± 95% CI of each sub-interval was quantified in embryos of the indicated genotypes. N is the number of embryos analyzed. **(B)** Stills of time-lapse movies of one-cell embryos of the indicated genotypes co-expressing LifeAct::GFP after mild depletion of NMY-2::mCherry^sen^. Selected time points show the actin equatorial band in the three situations. Statistical significance was determined using one-way ANOVA followed by Bonferroni’s multiple comparison test; ** P<0.001, *** P<0.0001, **** P<0.00001. Scale bar, 10 μm.

Next, we examined cortical myosin localization. NMY-2::mCherry^sen^ foci on the anterior cortex started to dissipate at anaphase onset in control embryos, and, coincident with actin equatorial enrichment, myosin patches appeared from the cytoplasm at the cell equator (Myosin AO-EA, 27±6 s). Myosin patches continued to accumulate at the equatorial region, resulting in the formation of a well-defined equatorial band 20 s later (Myosin EA-EB, 21±5 s) that was immediately followed by shallow deformation (Myosin EB-SD 11±3 s). In NMY-2(S251A) or NMY-2(R252A) embryos partially depleted of NMY-2::mCherry^sen^, myosin accumulation, like actin accumulation, was delayed relative to controls (Myosin AO-EA 75±26 s and 66±17s, respectively). Once an equatorial band of myosin was well-defined, deformation of the equator was significantly delayed (Myosin EB-SD 65±32 s and 52±19 s, respectively). In embryos partially depleted of NMY-2, myosin accumulation was very delayed relative to the accumulation of actin: first equatorial enrichment of myosin was observed 159±44 s after anaphase onset and the well-defined myosin equatorial band was observed 84 s later. In contrast to NMY-2(S251A) or NMY-2(R252A) embryos, once the myosin band formed, shallow deformation quickly followed (Myosin EB-SD 17±7 s; Supplemental Video 4; Fig. 7A). In sum, our data suggest that, independently of cortical flows, NMY-2 accumulation in the equatorial region is the rate limiting step for equatorial deformation, and that NMY-2 motor activity is essential for actin band compaction, proper alignment along the division plane and deformation of the equator.

## Discussion

### Myosin motor activity is required for cytokinesis

It is well established that myosin is essential for cytokinesis, but the importance of its motor activity, key to generating force and translocating actin filaments, remains unclear in animal cells. To investigate the role of myosin motor activity during cytokinesis, we generated and characterized *C. elegans* strains expressing two versions of motor-dead myosin, NMY-2(S251A) and NMY-2(R252A), and a partially motor-impaired myosin NMY-2(S250A). The equivalent mutations of NMY-2 S251A and R252A in other myosins consistently lead to inability of translocating actin filaments and abolishment of MgATPase activity (Shimada et al., 1997; Onishi et al., 1998a). According to detailed studies in *Dictyostelium*, the side chain of the serine directly coordinates to the magnesium ion in the ATPase pocket, and the side chain of the arginine forms a key salt bridge with the Switch II domain to stabilize a rotated state of the molecule that is required for ATP hydrolysis (Onishi et al., 1998a; b; Furch et al., 1999). Importantly, we show using co-sedimentation assays that both NMY-2(S251A) and NMY-2(R252A) are able to bind F-actin, and that they do so with similar (R252A) or even enhanced (S251A) affinity compared to their wild-type NMY-2. We also generated and characterized animals expressing equivalent mutations in the muscle myosin UNC-54. *Unc-54(S240A)* and *unc-54(R241A)* animals displayed a drastic reduction in motility and were unable to lay eggs, strongly suggesting that mutation of the highly conserved serine and arginine residue results in motor-dead myosin *in vivo*. Animals expressing UNC-54(S239A), which is equivalent to NMY-2(S250A), reduces but does not abolish motility, in agreement with the finding that this mutation partially compromises motor activity in *Dictyostelium* myosin (Shimada et al., 1997).

Our detailed analysis of embryos whose major source of NMY-2 is NMY-2(S251A) or NMY-2(R252A) demonstrates that myosin motor activity is essential for cytokinesis. *nmy-2(S251A)* or *nmy-2(R252A)* mutants only complete cytokinesis in the presence of transgene-encoded wild-type NMY-2 and do so with significantly slower kinetics than controls. This suggests that motor-dead NMY-2 hinders the activity of wild-type NMY-2, perhaps through the formation of heterotypic filaments like those made up of different isoforms of non-muscle myosin II or different classes of myosin (Beach et al., 2014; Shutova et al., 2014; Billington et al., 2015). Indeed, co-polymerization of different non-muscle myosin II isoforms has been shown to result in the formation of filaments with intermediate motile properties (Melli et al., 2017). Reinforcing the idea that myosin motor activity is essential for cytokinesis is the observation that embryos solely expressing the partially motor-impaired myosin NMY-2(S250A) delay in cytokinesis and are more sensitive to a reduction in overall myosin levels than wild-type embryos. This indicates that in the presence of motor-impaired myosin, more myosin molecules are needed to complete cytokinesis.

Our results contrast with the results of Ma et al., 2012 in COS-7 cells and mouse cardiomyocytes, where myosin was proposed to exert tension in a motor-independent manner during cytokinesis. However, we note that two of the three mutants used in this report, NMIIB(R709C) and NMIIA(N93K), should not be considered bona-fide motor-dead versions, as their characterization *in vitro* has been inconsistent. In fact, the mutant NMIIA(N93K) has very recently been clarified by the Adelstein and Sellers laboratories not to be a motor-dead myosin and to only present a modest decrease in enzymatic activity (Heissler et al., 2018). Also, the corresponding mutation of NMIIB(R709C) in NMIIA, R702A, was reported to translocate actin filaments at 50% the velocity of the wild-type counterpart (Hu et al., 2002). Finally, our *in vivo* characterization of NMY-2(R718C) and UNC-54(R710C), which are equivalent to NMIIB(R709C), strongly argues that this myosin mutant is not motor-dead, only motor-impaired. The third mutant used by Ma et al. is NMIIB(R234A), which is equivalent to NMY-2(R252A). NMIIB(R234A) is able to ameliorate cytokinesis defects in COS-7 cells depleted of NMIIB, while NMY-2(R252A) does not support cytokinesis in *C. elegans* embryos. This could indicate that cytokinesis operates differently in these two systems. Indeed, some adherent cultured cells can divide with a compromised contractile ring in an adhesion-dependent manner by migrating away in opposite directions during cell division (Kanada et al., 2005; Nagasaki et al., 2009; Dix et al., 2018).

### Myosin motor activity is required during furrow ingression

Embryos expressing motor-dead myosins NMY-2(S251A) or NMY-2(R252A) along with low levels of wild-type myosin (after RNAi of NMY-2::mCherry^sen^) were substantially delayed in contractile ring assembly, furrow initiation, and ring constriction. As the presence of motor-dead myosin led to actin bundle misalignment at the cell equator before furrow ingression, inappropriate F-actin architecture could contribute to ring constriction slowdown. However, embryos solely expressing partially motor-compromised NMY-2(S250A) or NMY-2(R718C) were able to assemble rings with normal timing and apparently normal F-actin alignment at the cell equator but were delayed in furrow initiation and ring constriction. This indicates that myosin motor activity continues to be required after ring formation and that motor activity is important to set the velocity of contraction of the ring’s F-actin network. Our results argue against the idea that the contractile stress driving ring constriction relies on passive actin filament cross linking combined with actin filament treadmilling, independent of myosin motor activity (Mendes Pinto et al., 2012; Oelz et al., 2015; Descovich et al., 2018).

It is possible that myosin motor activity contributes to ring constriction by determining actin filament turnover, as previously proposed (Guha et al., 2005; Murthy and Wadsworth, 2005; Wilson et al., 2010; Mendes Pinto et al., 2012). In agreement with data in budding yeast, when we partially deplete NMY-2, express NMY-2(S250A), or express a mix of NMY-2(S251A) or NMY-2(R252A) with wild-type NMY-2::mCherry^sen^, the reduction in ring constriction rate is accompanied by a proportional decrease in the total amount of actin in the ring, such that when actin levels are measured at 50% constriction they are similar to controls. This indicates that actin net depolymerization is also slowed down. Whether the simultaneous decrease in ring constriction rate and actin depolymerization is a direct effect of perturbing myosin remains to be determined. A direct effect could be due to decreased myosin-mediated actin filament buckling that likely contributes to net actin depolymerization. Actin filament buckling has been proposed to be caused by the action of myosin filaments with slightly different velocities (Lenz et al., 2012), and actin filament buckles have a higher curvature and should therefore be more prone to severing (Murrell and Gardel, 2012; Vogel et al., 2013). Even if the effect is direct, it is difficult to determine whether actin filament net depolymerization directly drives ring constriction or keeps ring structure optimized for myosin driven contractility.

### Cortical actin flows depend on myosin motor activity, but flows *per se* are not essential for ring formation or furrow ingression

It has been proposed that compressive persistent unidirectional flows toward the equator of the cell are the main driver for actin filament alignment at the equator and furrow formation in one-cell *C. elegans* embryos (Reymann et al., 2016). Alternatively, flow-independent recruitment of myosin to the cell equator could contribute to actin filament alignment and mediate ring formation and furrow ingression, as described in cultured cells (Zhou and Wang, 2008; Uehara et al., 2010; Spira et al., 2017). For analysis of cortical F-actin flows, we used embryos expressing LifeAct::GFP, which exhibits a cortical distribution profile that is identical to endogenously-tagged Plastin^PLST-1^::GFP and does not affect the kinetics of cytokinesis. By contrast, the previously used LifeAct::mKate2 probe exhibits a different cortical distribution profile from LifeAct::GFP and Plastin^PLST-1^::GFP, and we find that embryos expressing LifeAct::mKate2 complete cytokinesis with significantly slower kinetics. We confirm that robust cortical flows exist in embryos expressing LifeAct::GFP, but we find that flow intensity and direction is heavily influenced by the imaging method. The pronounced rotational flows observed under standard imaging conditions, in which embryos are subject to compression, are drastically decreased when embryos are imaged in an open chamber without compression. Importantly, imaging embryos without compression does not alter the kinetics of cytokinesis, suggesting that the importance of rotational flows for cytokinesis is minimal. Posterior-anterior directed flows in the posterior side of the embryo continue to be observed in these conditions and may contribute to the efficient alignment of actin filaments in the equatorial band. However, we find that even when unidirectional cortical flows are absent, as judged by PIV analysis in embryos co-expressing NMY-2(S251A) or NMY-2(R252A) and similar levels of wild-type NMY-2::mCherry, contractile rings form with normal timing, actin filaments align, and furrow ingression is successful (albeit slower because of the presence of motor-dead myosin). Overall, our results establish that cortical flows depend on myosin motor activity but argue against the idea that flows *per se* are critical for contractile ring assembly and constriction. It remains to be determined whether cortical flows are generated by myosin activity in the contractile ring that is able to pull cortical surface, as suggested by (Khaliullin et al., 2018), or whether myosin activity spread throughout the cortex potentiates flows in the direction of the furrow.

We propose that the rate limiting step for proper actin filament alignment at the division plane, compaction of the equatorial band, and deformation of the equator, is the accumulation of motor-competent myosin at the equatorial region, which may be largely independent of flows. This idea is supported by the observation that 1) when NMY-2 is substantially depleted, the remaining myosin is not used to flow actin filaments in the cortex, and actin filament band compaction and equatorial deformation occurs once enough myosin has been recruited to the equator; 2) in *nmy-2(S251A)* or *nmy-2(R252A)* embryos subject to RNAi to partially deplete NMY-2::mCherry^sen^, myosin and actin are recruited to the equator but the equatorial actin band takes longer to compact, and deformation of the equator is delayed. Myosin may be recruited to the equatorial region by a diffusion-equatorial retention mechanism, whereby myosin that will form the contractile ring diffuses from the cytoplasm and is retained at the equator due to myosin filament formation mediated by the phosphorylation of its regulatory light chain by kinases that are locally activated by Rho-GTP (Kimura et al., 1996; Bement et al., 2005; Uehara et al., 2010). Equatorially-localized myosin motor activity then drives the alignment and compaction of actin filament bundles, likely aided by actin cortical flows, to bring about equatorial deformation and furrow ingression.

## Material and methods

### *C. elegans* strains

Strains used in this study are listed in Table S1 and were maintained on nematode growth medium (NGM) plates seeded with OP50 *E. coli* at 20°C.

Re-encoded nmy-2 transgene fused to mCherry in strain GCP22 was generated by Mos1 mediated single copy transgene insertion (MosSCI) on chromosome II, tTi5605 (Frøkjaer-Jensen et al., 2008; Frøkjær-Jensen et al., 2012). A region of ∼400 bp in exon 12 was re-encoded so that transgenic nmy-2, and not endogenous nmy-2, could be specifically depleted by RNAi. Single copy transgene insertions were generated by injecting a mixture of target plasmid, transposase plasmid pCFJ601, and selection markers into the strain EG6429 as described previously (Frøkjaer-Jensen et al., 2008; Frøkjær-Jensen et al., 2012). Transgene integrations were confirmed by PCR of regions spanning each side of the insertion, sequencing of the entire genomic DNA locus and by fluorescence microscopy. The endogenous loci of *nmy-2* and *unc-54* were modified using the CRISPR-Cas9 technique for mutation insertion. Two or three single guide RNAs (sgRNAs) were cloned into the pDD162 vector (Dickinson et al., 2013) and injected together with a single stranded repair template carrying the modified sequence of interest flanked by 35-50 bp homology regions (IDT ultramer) in the gonads of young adult worms. To facilitate the identification of successfully injected animals, injection mixes contained a sgRNA and a repair template to insert the R92C mutation in the dpy-10 gene, which causes a dominant roller phenotype in modified animals (Levy et al., 1993; Arribere et al., 2014). All point mutations were verified by genomic DNA sequencing.

### RNA interference

RNAi was performed by feeding worms with bacteria that expressed the dsRNA of interest. DNA fragments of interest were cloned into the L4440 vector and these were transformed into HT115 *E. coli,* which were used to feed worms after IPTG induction. Primers in table S2 were used to amplify nmy-2 DNA fragments from N2 cDNA (nmy-2_RNAi#1) or from a synthesized re-encoded region (nmy-2_RNAi#2). These fragments were cloned into the EcoRV site in the L4440 vector. For feeding RNAi of *myo-3*, *unc-54* or *par-2*, corresponding L4440 vectors were obtained from the Ahringer library (Source Bioscience) and sequenced to confirm gene target. In *nmy-2*(*RNAi*) experiments, depletion of endogenous NMY-2 using NMY-2_RNAi#1 was performed in L4 worms at 20 °C. For partial depletion, GCP21 or GCP179 worms were incubated for 34-40 hours (Fig. 5, 7A, S3, S5A, S5E). For mild depletions, GCP21 and GCP401 worms were incubated for 25–28 hours (Fig. 4H, J). Depletion of transgenic NMY-2::mCherry^sen^ using nmy-2_RNAi#2 was performed in GCP22, GCP618 or GCP592 L4 worms at 20 °C. For mild depletions, worms were incubated for 22-26 hours (Fig. 2D, 3B, 3C, 5A, 7A-B, S4C-D, S5A, S5E). For penetrant depletion, worms were incubated for 44-48 hours (Fig. 2D). Depletion of MYO-3 was accomplished by incubating L1 worms of strains GCP565 and GCP619 (Fig. 1D), GCP523 (Fig. 4A) and GCP524 (Fig. S2B) for 72 hours at 20°C. Depletion of MYO-3, UNC-54 or simultaneous depletion of MYO-3 and UNC-54 in figure S1B were performed by feeding L1 worms of N2 strain for 72 hours at 20 °C. For double RNAi of *myo-3* and *unc-54*, bacteria expressing each of the constructs were mixed 1:1 before seeding the RNAi plates (Fig. S1B). Depletion of *par-2* was performed in GCP21 and GCP22 L4 worms for 48-52 hours at 20 °C (Fig. S4C-D).

### Live imaging

Gravid hermaphrodites were dissected and one-cell *C. elegans* embryos were mounted on a drop of M9 (86 mM NaCl, 42 mM Na_2_HPO_4_, 22 mM KH_2_PO_4_, and 1 mM MgSO_4_) in 2% agarose pads overlaid with a coverslip. In this conventional manner of imaging *C. elegans* embryos, they are subject to compression. In figure S5B-D embryos were placed on imaging chambers (Carvalho et al., 2011) where they are immersed in M9 and not subject to compression. Live imaging of cytokinesis was performed at 20 °C. Images were acquired on a spinning disk confocal system (Andor Revolution XD Confocal System; Andor Technology) with a confocal scanner unit (CSU-22; Yokogawa) mounted on an inverted microscope (Ti-E, Nikon) equipped with a 60 ×; 1.42 oil-immersion Plan-Apochromat objective, and solid-state lasers of 488 nm (250 mW) and 561 nm (250 mW). For image acquisition an electron multiplication back-thinned charge-coupled device camera (iXon; Andor Technology) was used. Acquisition parameters, shutters, and focus were controlled by Andor iQ3 software. For central plane imaging in one-cell embryos, 6 ×; 1–μm z stacks were collected in the 488-nm and 561-nm channels every 10 seconds through the center of the embryo. For cortical imaging in one-cell embryos, 7 ×; 0.5–μm z stacks were collected in the 488-nm channel every 5 seconds. For cortical flow imaging in one-cell embryos, 7 ×; 0.5–μm z stacks were acquired every 5 seconds in embryos of GCP22, GCP401, GCP592 and GCP618 strains. For liquid thrashing assays, N2, GCP523, GCP524, GCP565 or GCP619 worms were synchronized at the L1 stage by performing alkaline bleach treatment of adult worms (0.8% bleach, 250 mM NaOH; Stiernagle, 2006) to extract embryos, which were left to hatch overnight in M9 medium. L1s were then plated and grown at 20 °C. Young adult worms were transferred to an M9 droplet and left to acclimatize for 2 min after which images were acquired at ∼40 fps average.

### Calculation of embryonic viability percentage and egg laying rate

In figures 4G and S2D, L4 worms of strains GCP21, GCP401 or GCP420 were placed on NGM plates and singled out onto fresh plates after 40 hours at 20°C. Worms were let to lay eggs for eight hours. In figure 2D, GCP22, GCP618 or GCP592 L4 worms were placed on plates with bacteria expressing nmy-2_RNAi #2 for 22 hours (mild depletion) or 40 hours (penetrant depletion) at 20 °C. Worms were then singled out onto fresh RNAi plates and let to lay eggs for 4 hours (mild depletion) or 8 hours (penetrant depletion). In all cases mother worms were removed after the egg laying interval and embryos were left to hatch at 20°C for 24 hours before counting. The number of hatched and unhatched (dead) embryos was counted and the percent embryonic viability was calculated by dividing the number of hatched embryos by the total number of progeny. To measure egg laying rates in figures 1E, 4B and S2C, L4 worms of strains N2, GCP523, GCP524, GCP565 or GCP619 were placed on NGM plates and singled out onto fresh plates after 40 hours at 20°C. Worms were let to lay eggs for eight hours. The rate was calculated by dividing the total number of eggs by the number of laying hours.

### Preparation of worm protein extracts and immunoblotting

For figures 2C and 4F, sixty L4 worms of each of the strains: N2, GCP22, GCP592 GCP618, or GCP401 were grown on NGM plates for 46 hours at 20 °C. For protein sample preparation, worms were collected and washed three times in M9 medium with 0.1% triton X-100 and pelleted at 750 xg. Worm lysis was performed by resuspending the worm pellet in 1x Laemmli buffer with ⅓ volume of quartz sand (Sigma). Tubes were subject to three 5-minute cycles of alternating boiling at 95 °C and vortexing after which the quartz sand was pelleted and the supernatant recovered. Protein samples were resolved by SDS-PAGE in a 7.5% acrylamide gel and transferred to 0.2-μm nitrocellulose membranes (GE Healthcare). Membranes were blocked with 5% non-fat dry milk in TBST (20 mM Tris, 140 mM NaCl, and 0.1% Tween, pH 7.6) and probed at 4°C overnight with the following primary antibodies: anti-NMY-2 antibody, 1:10000 (rabbit polyclonal against residues 945-1368); anti-tubulin, 1:5000 (mouse monoclonal DM1-α, Sigma). Membranes were washed three times with TBS-T, incubated with HRP-conjugated secondary antibodies donkey anti-rabbit 1:5000 or goat anti-mouse 1:5000 (Jackson ImmunoResearch) for 1 hour at room temperature, and washed again three times with TBST. Blots were visualized by chemiluminescence using Pierce ECL Western Blotting Substrate (Thermo Fisher Scientific) and imaged in a ChemiDoc™ XRS+ System with Image Lab™ Software (Bio-Rad).

### Protein alignments

Myosin protein sequences for alignments shown in figures 1B, 1C, S1A, S2A were obtained from the Uniprot database with the following accession numbers: P08799|MYS2_DICDI; G5EBY3|G5EBY3_CAEEL; Q20641|Q20641_CAEELQ99323|MYSN_DROME; F1QC64|F1QC64_DANRE; F8W3L6|F8W3L6_DANRE; F1R3G4|F1R3G4_DANRE; 93522|O93522_XENLA; Q04834|Q04834_XENLA; Q8VDD5|MYH9_MOUSE; Q61879|MYH10_MOUSE; Q6URW6|MYH14_MOUSE; P35579|MYH9_HUMAN; P35580|MYH10_HUMAN; Q7Z406|MYH14_HUMAN; P02566|MYO4_CAEEL and aligned using the Jalview software (Waterhouse et al., 2009) and the muscle algorithm with default parameters (Edgar, 2004).

### NMY-2 S1 fragment expression and purification

cDNA sequences of wild-type or mutant NMY-2 S1 fragments (amino acids 1-874), full length MLC-4 and MLC-5 were cloned in the pACEbac1 expression vector. NMY-2 constructs were tagged N-terminally with 6×;His followed by a linker. Myosin light chains MLC-4 and MLC-5 were tagged N-terminally with a Strep-tag II followed by a linker. Bacmid recombination and virus production were performed as described previously (Bieniossek et al., 2008). NMY-2 S1, MLC-4 and MLC-5 baculovirus were used to co-infect 500-mL cultures of Sf21 cells (0.8 ×; 10^6^ cells/ml, SFM4 medium; Hyclone). Cells were harvested by centrifugation at 3000 xg for 5 min. Pellets were resuspended in lysis buffer (15 mM MOPS, 300 mM NaCl, 15 mM MgCl2, 0.1% Tween 20, 1 mM EDTA, 3 mM NaN_3_, 1 mM DTT, pH 7.3) supplemented with EDTA-free Complete Protease Inhibitor Cocktail (Roche), sonicated, and incubated for 20 minutes with 1 mM ATP to detach myosin from actin. Lysates were then cleared by centrifugation at 34000 xg for 40 minutes. The complex was purified by batch affinity chromatography using Strep-Tactin Sepharose (IBA). Beads were washed with wash buffer (10 mM MOPS, 500 mM NaCl, 5 mM MgCl2, 0.1% Tween 20, 1 mM EDTA, 3 mM NaN_3_, 1 mM DTT pH 7.3) supplemented with 1 mM ATP in the first wash and eluted on a gravity column with elution buffer (10 mM MOPS, 500 mM NaCl, 2.5 mM Desthiobiotin, 3 mM NaN_3_, 1 mM DTT, pH 8.0). The complex was further purified by size-exclusion chromatography using a Superose 6 increase 10/300 column (GE HealthCare) pre-equilibrated with 10 mM MOPS, 500 mM NaCl, 1 mM EDTA, pH 7.3. Fractions containing the S1 myosin complex were pooled, and glycerol and DTT were added to a final concentration of 10% (vol/vol) and 1 mM, respectively. Aliquots were flash-frozen in liquid nitrogen and stored at −80°C.

### F-actin co-sedimentation assays

Wild-type or mutant myosin S1 complex were dialysed overnight in actin buffer (5 mM Tris, 0.2 mM CaCl_2_, 50 mM KCl, 2 mM MgCl_2_, 1 mM DTT, pH=8). All proteins were pre-cleaned by centrifugation at 150000 xg for 1 hour in an Optima XP centrifuge with a TLA-100 rotor (Beckman-Coulter) prior to assays. F-actin was prepared using the *Non-muscle actin binding protein spin-down* assay kit (Cat. # BK013 Cytoskeleton Inc.) according to manufacturer’s instructions. Briefly, human platelet actin was polymerized in actin polymerization buffer (5 mM Tris, 0.2 mM CaCl_2_, 50 mM KCl, 2 mM MgCl_2_, 1 mM ATP) to obtain a 21 μM F-actin stock. For the co-sedimentation assays, an equal amount of each myosin complex was incubated with 14.7 μM F-actin (final concentration) or a similar volume of buffer (as negative control), for 30 minutes. Samples were then centrifuged at 150000 xg for 1.5 hours and supernatants were carefully removed and Laemmli sample buffer was added to 1x final concentration. To resuspend the pellet fractions, 30 μL of water were added to the tubes and mixed by pipetting every 2 minutes over a period of 10 minutes on sice. Laemmli sample buffer was added to pellet fractions to obtain 1x final concentration. For coomassie stained gels in figures 1F, 4C, S2B, Samples were incubated for 4 min at 95 °C to denature proteins and run on 20-4% TGX gradient precast gels (Bio-Rad) using Tris-Glycine-SDS running buffer.

### Body wall muscle actin staining

Phalloidin labelling of muscle actin shown in figure S1C in N2, GCP523, GCP565 and GCP619 worms was performed as previously reported (Ono, 2001). Briefly, adult worms were collected and washed twice in M9 buffer, fixed with 4% formaldehyde in 1x cytoskeleton buffer (10 mM MES-KOH, pH 6.1, 138 mM KCl, 3 mM MgCl_2_, 2 mM EGTA) containing 0.32 M sucrose (Cramer and Mitchison, 1993) for 15 min, permeabilized with acetone at −20 °C for 5 min, washed with PBS containing 0.5% Triton X-100 and 30 mM glycine (PBS-TG) for 10 min, and stained with 1:40 Oregon Green phalloidin (Molecular Probes, Thermo Fisher Scientific) in PBS-TG for 30 min. After three 10-minute washes with PBS-TG, worms were mounted with ProLong Antifade containing DAPI (Molecular Probes, Thermo Fisher Scientific). Processing of N2, GCP513 and GCP623 worms in figure 2A was done as described above but only DAPI staining of the gonads is shown.

### Image analysis, quantifications, and statistics

All microscopy image processing and measurements were done using Fiji (ImageJ; National Institutes of Health; (Schindelin et al., 2012) and Matlab (MathWorks). Z-stacks taken on the cell cortex were projected using the maximum intensity projection tool. Images within each panel were scaled equally. The equatorial region of the central plane was selected to create the kymographs shown in figures 3B, 4H, S3A’, using the Make Montage tool. Flow kymographs shown in figures 6A’ and S5A were created using the Dynamic Reslice tool after tracing a 100 pixel-long vertical line on the anterior region of the cortical plane as illustrated in figure 6A. Time projection in figure S5B was done using the Temporal color code tool in the maximum projected image of the cortical region of a control embryo expressing LifeAct::GFP. Coomassie stained SDS-PAGE gels were digitized in a GS-800 Calibrated Imaging Densitometer (Bio-Rad) and relative band intensity quantified in image-Lab 5.2.1 (Bio-Rad). For each myosin mutant, the ability of myosin to co-sediment with F-actin was quantified by dividing the intensity of the band corresponding to NMY-2 S1 fragment in the pellet by the sum of intensities of the NMY-2 S1 fragment bands in the supernatant and pellet. The average of at least 2 independent experiments was used in the graphs of figures 1G, 4D, S2F. Graph plotting, linear regressions and statistical analyses were performed with Prism 6.0 (GraphPad Software). All error bars represent the 95% CI of the mean and statistical significance tests were performed using Student-t or one-way ANOVA with Bonferroni correction as indicated in figure legends.

### Measurement of ring assembly and furrow initiation time intervals, and ring constriction rate

Measurement of ring assembly and furrow initiation time intervals, as well as overall ring constriction rate were performed during cytokinesis in one-cell embryos of the following strains: GCP22, GCP592 and GCP618 (Fig. 3C); GCP21 and GCP401 (Fig. 4I-J); GCP21 and CGP420 (Fig. S2G); GCP179 (Fig. S3A); OD121, GCP113, SWG001 and RZB213 (Fig. S4A); GCP21 (Fig. S5D). This analysis only included embryos that completed ring constriction. Assembly time was the time interval between anaphase onset and the establishment of a shallow deformation in the equatorial region. Furrow initiation corresponded to the time interval between the establishment of the shallow deformation and the time when the plasma membranes of the nascent daughter cells became juxtaposed to one another (back-to-back membrane configuration). During ring constriction, the distance between the two sides of the ring on the z-plane where this was the widest was measured for each time point and plotted against time. Ring constriction rate was the slope of the linear region between ∼70% and 30% ingression. All these intervals were determined based on imaging of the embryo central plane.

### Fluorescence intensity measurements

To quantify the levels of Lifeact::GFP at the tip of the cytokinetic furrow in one-cell embryos, the mean fluorescence intensity in a 10-pixel wide, 30-pixel long (1.8×5.3 μm) box drawn over the tip of the furrow at 50% ingression was determined and the mean camera background was subtracted (Fig. 5A). In figure 5A the average intensity at the furrow tip is presented as a percentage of the corresponding controls. In figure 5B, quantification of actin levels in the contractile ring of ABa cells was done in GCP22 4-cell embryos by manually tracing a 0.7 μm-thick line along the entire ring. The mean GFP fluorescence intensity along the 0.7 μm-line was determined for each time point and the mean fluorescence intensity in a circle drawn over the cytoplasm for each time point was subtracted. Before quantification, each time-lapse movie was corrected for the intensity decay using the imageJ tool bleach correction and the method simple ratio. Data from multiple rings were pooled and plotted against ring perimeter. The mean of data points that fell in overlapping 5-μm intervals was calculated and plotted against the perimeter at the center of each interval.

### Measurement of body bends swimming frequency

Body bends swimming frequencies in figures 1D, 4A, S2B were automatically quantified using the wrMTrck plugin with standard parameters (Nussbaum-Krammer et al., 2015) in Fiji. Image background was removed by subtracting the average intensity projection of the stack and worms were segmented using Otsu intensity thresholding.

### Measurement of time required for actin and myosin equatorial appearance and band formation

In figures 7A and S4D, the time for cortical actin or myosin equatorial appearance (EA) was determined as the first time point after anaphase onset when signal of actin/myosin were observed in the equatorial region, whereas the actin or myosin equatorial band (EB) corresponded to the time point when a band of filaments/patches was observed across the whole equator. The time of anaphase onset and shallow deformation was determined based on imaging of the central plane of the embryo.

### Analysis of cortical flows

Cortical flow dynamics was analyzed in one-cell embryos of the strains GCP21, GCP401, GCP562, and GCP618 between anaphase onset and back to back membrane configuration. In figures 6B’, 6C, S5C and S5E, magnitude and direction of cortical flow were quantified using an Iterative Particle Image Velocimetry (PIV) plugin for ImageJ (Tseng et al., 2012). A 192 128 pixel (34.13 ×; 22.76 μm) region of interest was applied to all embryos to standardize the cortex area and exclude cortical curved peripheries. To average the maximum particle displacement with better resolution, two iterations were performed, in which the application of a larger interrogation window of 64 ×; 64 pixel (first iteration) was followed by a smaller interrogation window of 32 ×; 32 pixel (second iteration). The interrogation windows did not overlap each other. The intensity and direction of the flow field shown in the magnitude-arrow plots in figure 6B’ were generated using the plot function of the plugin. After flow field generation, the 192 ×; 128 pixel region was divided in two adjacent sub-regions along anterior-posterior axis of the embryo for quantification: anterior cortex (1st-108th pixel; 0.17-19.2 μm) and posterior cortex (109th-192nd pixel; 19.38-34.13 μm). The furrow region corresponded to part of each of the two sub-regions (92nd-120th pixel; 16.36-21.33 μm). Each velocity vector was resolved into components along the anterior-posterior axis, Vx, and perpendicular to the anterior-posterior axis, Vy. The mean Vx and Vy were estimated for each time point and plotted as a function of time to trace the cortical flow profiles in figures 6C, S5C and S5E.

## Acknowledgments

We would like to acknowledge Stephan Grill, Ronen Zaidel-Bar and Karen Oegema for providing strains for this study and Claudia Brito and Sandra Sousa for advice with myosin biochemistry. The research leading to these results has received funding from the European Research Council under the European Union’s Horizon 2020 research and innovation programme to AXC (grant agreement 640553 - ACTOMYO), and under the European Union’s Seventh Framework programme (FP7/2007-2013) to RG (grant agreement ERC-2013-StG-338410-DYNEINOME), national funds through Fundação para a Ciência e a Tecnologia (FCT) under the project FCOMP-01-0124-FEDER-028255 (PTDC/BEX-BCM/0654/2012) and from Norte-01-0145-FEDER-000029 - Advancing Cancer Research: From basic knowledge to application, supported by Norte Portugal Regional Operational Programme (NORTE 2020), under the PORTUGAL 2020 Partnership Agreement, through the European Regional Development Fund (FEDER). AXC and RG have FCT Investigator positions funded by FCT and co-funded by the European Social Fund through Programa Operacional Temático Potencial Type 4.2 promotion of scientific employment. FYC and AMS hold FCT post-doctoral fellowships SFRH/BPD/93528/2013 and SFRH/BPD/95707/2013, respectively. AFS holds an FCT PhD scholarship SFRH/BD/121874/2016. DSO also received funding from the Programa Operacional Regional do Norte under the Quadro Estratégico Nacional through FEDER and by FCT grant NORTE-07-0124-FEDER-000003 (Cell Homeostasis Tissue Organization and Organism Biology). The authors declare no competing financial interests. The funders had no role in study design, data collection and analysis, decision to publish, or preparation of the manuscript.

## Author contributions

AXC, RG, DO conceived and designed experiments. AXC, RG, DO wrote the manuscript. DO, FYC, JS, JL, AXC prepared the figures and movies. DO, FYC, JS, JL, FS, AMS performed the experimental work and analyzed the data.

